# Inter-session repeatability of markerless motion capture gait kinematics

**DOI:** 10.1101/2020.06.23.155358

**Authors:** Robert M. Kanko, Elise Laende, W. Scott Selbie, Kevin J. Deluzio

## Abstract

The clinical uptake and influence of gait analysis has been hindered by inherent flaws of marker-based motion capture systems, which have long been the standard method for the collection of gait data including kinematics. Markerless motion capture offers an alternative method for the collection of gait kinematics that presents several practical benefits over marker-based systems. This work aimed to determine the reliability of lower limb gait kinematics from video based markerless motion capture using an established experimental protocol for testing reliability. Eight healthy adult participants performed three sessions of five over-ground walking trials in their own self-selected clothing, separated by an average of 8.5 days, while eight synchronized and calibrated cameras recorded video. 3D pose estimates from the video data were used to compute lower limb joint angles. Inter-session variability, inter-trial variability, and the variability ratio were used to assess the reliability of the gait kinematics. Relative to repeatability studies based on marker-based motion capture, inter-trial variability was slightly greater than previously reported for some angles, with an average across all joint angles of 2.2°. Inter-session variability was smaller on average than previously reported, with an average across all joint angles of 2.5°. Variability ratios were all smaller than those previously reported with an average of 1.2, indicating that the multi-session protocol increased the total variability of joint angles by only 20% of the inter-trial variability. These results indicate that gait kinematics measured using markerless tracking were less affected by multi-session protocols compared to marker-based motion capture.

## 1. Introduction

Three-dimensional (3D) human movement analysis is a widely used tool in clinical and research biomechanics to provide comprehensive 3D representations and quantification of individuals’ movement patterns, particularly gait. This tool allows comparisons to be made within and between individuals and groups on a singular or longitudinal basis, providing quantified measures of functional musculoskeletal health. This data has the potential to help monitor disease progression, improve physical therapy, and aid in surgical decision-making (Astephen et al., 2008; Crowell and Davis, 2011; Haim et al., 2012; Smania et al., 2011; Wren et al., 2011). However, the clinical uptake and influence of gait analysis has been hindered by technological limitations of marker-based motion capture.

Marker-based motion capture has several inherent flaws that affect a researcher or clinician’s ability to collect repeatable measures of gait, particularly over multiple visits. These limitations include lengthy data collection times, the requirement of highly skilled experimenter capable of accurate marker placement, dedicated laboratory space, and the intrusive nature of the technology and experimental protocols on the subject or patient, which in turn limit the use of this data in clinical decision-making (Narayanan, 2007; Benedetti et al., 2017). This technology requires markers to be placed on patients’ palpable anatomical landmarks, a process that requires 20-30 minutes and causes the resulting data to be susceptible to inaccurate and inconsistent marker placement (Della Croce et al., 2005). The issues associated with marker placement are especially evident in data collected by different operators, during separate collection sessions, and between laboratories (Gorton et al., 2009). The changing collection conditions can lead to inconsistent findings or conclusions across gait studies, demonstrating that clinically acceptable errors are possible but not always achieved in gait analysis (McGinley et al., 2009). Standardized protocols have been shown to reduce undue variability in data collected on the same subject at different laboratories by different operators (Gorton et al., 2009); however, some inter-operator variability persists even when an identical collection protocol and the same laboratory are used (Maynard et al., 2003). While effective alternative anatomical calibration techniques have been demonstrated that do not require highly trained operators, they are still time intensive and somewhat intrusive (Donati et al., 2008). Marker-based motion capture technology requires a dedicated laboratory space or enclosed environment, which is a significant barrier to the collection of gait data outside of research facilities and may be a foreign environment to subjects. Finally, subjects are required to wear minimal, skin-tight clothing so as to enable researchers or clinicians to attach markers to their body. In combination, the unfamiliar laboratory environment, physical and social discomfort due to the clothing and markers, and requests to perform natural gait on demand reduce the fidelity of gait data collected using marker-based motion capture and negatively affect recruitment rates.

Markerless motion capture is a quickly evolving technology that offers an alternative to measure human movement without the need for skin-based markers or sensors. These systems often use arrays of two-dimensional (2D) video cameras or depth sensors in combination with machine learning algorithms to estimate human pose during physical tasks and have been implemented to varying levels of success (Mathis et al., 2018; Mathis et al., 2020). With recent advances in computer vision techniques and the increased availability of computational power, markerless motion capture systems have undergone significant improvements in processing time and accuracy and are now available as commercial products. *Theia3D* (Theia Markerless Inc., Kingston, ON) is one example of a machine learning-based markerless motion capture software that uses 2D video data from an array of standard video cameras to perform 3D pose estimation on human subjects. Since the motion capture system does not rely on skin-based markers, subjects are not required to wear minimal and skin-tight clothing, and instead wear their own clothing. Thus, subjects may be significantly more comfortable and able to perform physical tasks more naturally, leading to more ecological data. Furthermore, the markerless system is not limited to use in laboratory spaces, allowing data to be collected in real-world environments which cannot be replicated in the laboratory. Finally, since the markerless system is not reliant on markers that can be placed incorrectly or inconsistently, there may be less variability introduced to kinematic measurements across sessions or operators.

The objective of this work was to determine the reliability, in the form of test-retest repeatability, of over-ground gait kinematics measured using the *Theia3D* markerless motion capture system and compare those to previously reported values for field-accepted marker-based motion capture systems. We hypothesized that the markerless motion capture system would have lower variability in joint kinematics between repeated visits compared to marker-based systems.

## 2. Methods

### 2.1 Theia3D Markerless Motion Capture

*Theia3D* is a deep learning algorithm-based approach to markerless motion capture which uses deep convolutional neural networks for object recognition (humans and human segments) within 2D camera views (Mathis et al., 2020). The neural networks were trained on over 500,000 images sourced from Microsoft COCO (Lin et al., 2015) and a proprietary set of images. A variety of joint locations and other identifiable anatomical features in the images were manually labelled by highly trained annotators and controlled for quality by a minimum of one additional expert labeller. These training images consisted of humans in a wide array of settings, clothing, and performing various activities. Deconvolutional layers are used to produce spatial probability densities for each image, representing the likelihood that an anatomical feature is in a particular location. During training with labeled data, the weights are iteratively adjusted. For a given image, the network assigns high probabilities to labeled anatomical feature locations and low probabilities elsewhere. This learning that occurs during training enables the application of “rules” for identifying the learned features within a new image.

When using *Theia3D* for markerless motion capture, the user provides newly collected video data from multiple synchronized and calibrated video cameras that capture one or more subjects performing a physical task. The time required to collect data is largely dependent on the task of interest but can take less than five minutes for the collection of ten walking trials, for example. From the collected videos, *Theia3D* extracts the 2D positions of its learned features within all frames of all of the videos, which are then transformed to 3D space based on the computed position and orientation of the cameras. Finally, an articulated multi-body model is scaled to fit the subject-specific landmarks positions in 3D space, and a multi-body optimization approach (inverse kinematic (IK)) is used to estimate the 3D pose of the subject throughout the physical task. By default, the lower body kinematic chain has six degrees-of-freedom (DOF) at the pelvis, three DOF at the hip, two DOF at the knee (flexion/extension and ab/adduction), and three DOF at the ankle. If so inclined, the user may opt to further constrain the knee to only one DOF or remove the translation constraints to allow six DOF (6DOF) at the ankle.

### 2.2 Participants

A convenience sample of eight healthy, recreationally active adults (2 female, mean (SD) age: 30.3 (14.1) years, height: 173.8 (9.0) cm, mass: 69.0 (12.4) kg) were recruited to participate in this multisession study at the Human Mobility Research Laboratory (Kingston, ON). Participants gave written informed consent, and this study was approved by the institutional ethics board. Exclusion criteria included having any neuromuscular or musculoskeletal impairments that could prevent their performance of walking. Participants were given no prior instruction for what clothing to wear and participated wearing the clothing in which they arrived, and either their personal running shoes or were provided with a pair of running shoes. Participants returned for a total of three sessions, which were separated by an average of 8.5 (2.0) days. A composite image of the clothing worn by participants during each session is shown in Figure 1.

**Figure 1:**
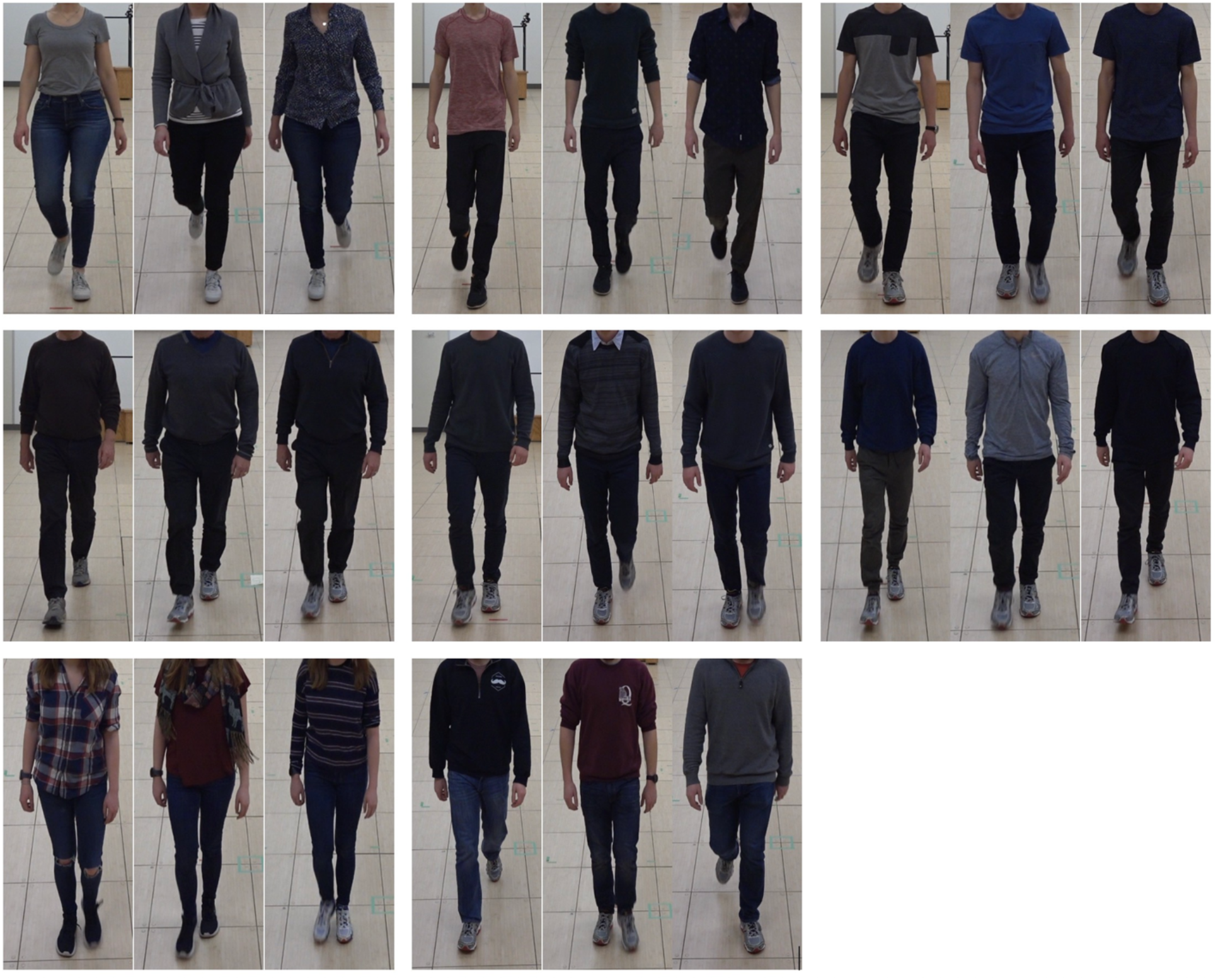
Composite image of participants and their clothing, with three images per participant (one per session). Participants were given no specific instructions regarding the clothing they should wear during the data collections.

### 2.3 Experimental Setup and Data Collection Procedure

Eight Sony RX0 II cameras (Sony Corporation, Minato, Japan) were connected and synchronized using a Sony Camera Control Box and were arranged around a capture volume approximately 12 metres long by 5 metres wide by 2.5 metres tall within a large indoor laboratory space. The camera positions were minimally repositioned between the collection sessions and were recalibrated before each session. Red tape lines were placed on the ground ten metres apart and were used as walkway start/finish lines. At every session, participants performed five over-ground walking trials between the red start/finish lines at their comfortable walking speed, alternating direction for each trial, while synchronized 2D video data were collected at 60 Hz.

### 2.5 Data Analysis

Video data were processed twice using *Theia3D* software to obtain 3D pose estimates of the subjects using the standard IK solution and the 6DOF ankle option. In both cases, the 3D pose estimates for each segment of the articulated multi-body model were exported as 4×4 pose matrices for each frame of data for further analysis in Visual3D (C-Motion Inc., Germantown, MD). A built-in Visual3D model designed for use with *Theia3D* output data was applied to all trials, and virtual toe and heel markers were added to the model based on *Theia3D* estimates of their location. These virtual markers were used with the method described by Zeni *et al.* (Zeni et al., 2008) to determine heel-strike and toe-off gait events throughout each trial. Lower limb joint angles were calculated using the standard Cardan rotation sequence (X-Y-Z), equivalent to the joint coordinate system (Grood and Suntay, 1983), and time-normalized to the gait cycle using the heel-strike events. Using the first left and right gait cycle that occurred within the central five-meter portion of the walkway, average time-normalized joint angles were obtained for each trial and each joint and were exported for further analysis in MATLAB (The MathWorks Inc., Natick, MA).

Repeatability of measured kinematics was assessed using the method described by Schwartz *et al.* which measures subjects’ inter-session variability, inter-trial variability, and the ratio between them (Schwartz et al., 2004). The inter-trial variability captures the stride-to-stride variation that exists within each kinematic measure due to intrinsic subject variability including the effects of varying gait speed, and any systematic noise that may be present between successive trials. The inter-session variability captures any variation that arises due to the repeated sessions methodology in addition to inter-trial variation. The variability ratio gives the proportion of inter-session variability that is accounted for by the inter-trial variability. The term “variability” is used in this work in place of “error”, as used by Schwartz *et al.* when describing the measure obtained from the standard deviation of the inter-trial, inter-session, and interoperator deviations, as error implies a difference from a ground truth measurement, which in general cannot be assessed for motion capture systems. These repeatability measures were compared to those in existing literature that utilized marker-based motion capture systems and the Schwartz *et al.* method (Caravaggi et al., 2011; Kaufman et al., 2016; Manca et al., 2010; Schwartz et al., 2004). Among these studies, one used a modified version of the Conventional Gait Model (a direct kinematic (DK) approach) (Schwartz el al., 2004) and three used 6DOF biomechanical models (Manca et al., 2010; Caravaggi et al., 2011; Kaufman et al., 2016). Since IK techniques have been shown to reduce measurement variability (Charlton et al., 2004; Kainz et al., 2017), we performed our analyses using both the default IK model and the optional 6DOF ankle model implemented in *Theia3D*.

## 3. Results

Using markerless motion capture, the time required for each session, including subject initiation and data collection, was typically between five and ten minutes. The average standard deviation in gait speed for all eight subjects across all three collection sessions was 0.054 m/s, indicating their self-selected over-ground walking speed varied little across sessions.

Joint angle waveforms for one representative subject are presented in Figure 2. The variability seen in each joint angle in Figure 2 is reflected in the inter-trial variability, inter-session variability, and variability ratio measures shown in Figure 3, and summarized using the average across all subjects in Figure 4. Repeatability measures were found to differ minimally between the kinematics obtained from *Theia3D* using the IK model and the 6DOF ankle model, with the greatest difference being a decrease of 0.2° in both the inter-trial and inter-session variability of ankle internal/external rotation for the IK model compared to the 6DOF ankle model. Subsequent results mentioned will be those of the IK model.

**Figure 2:**
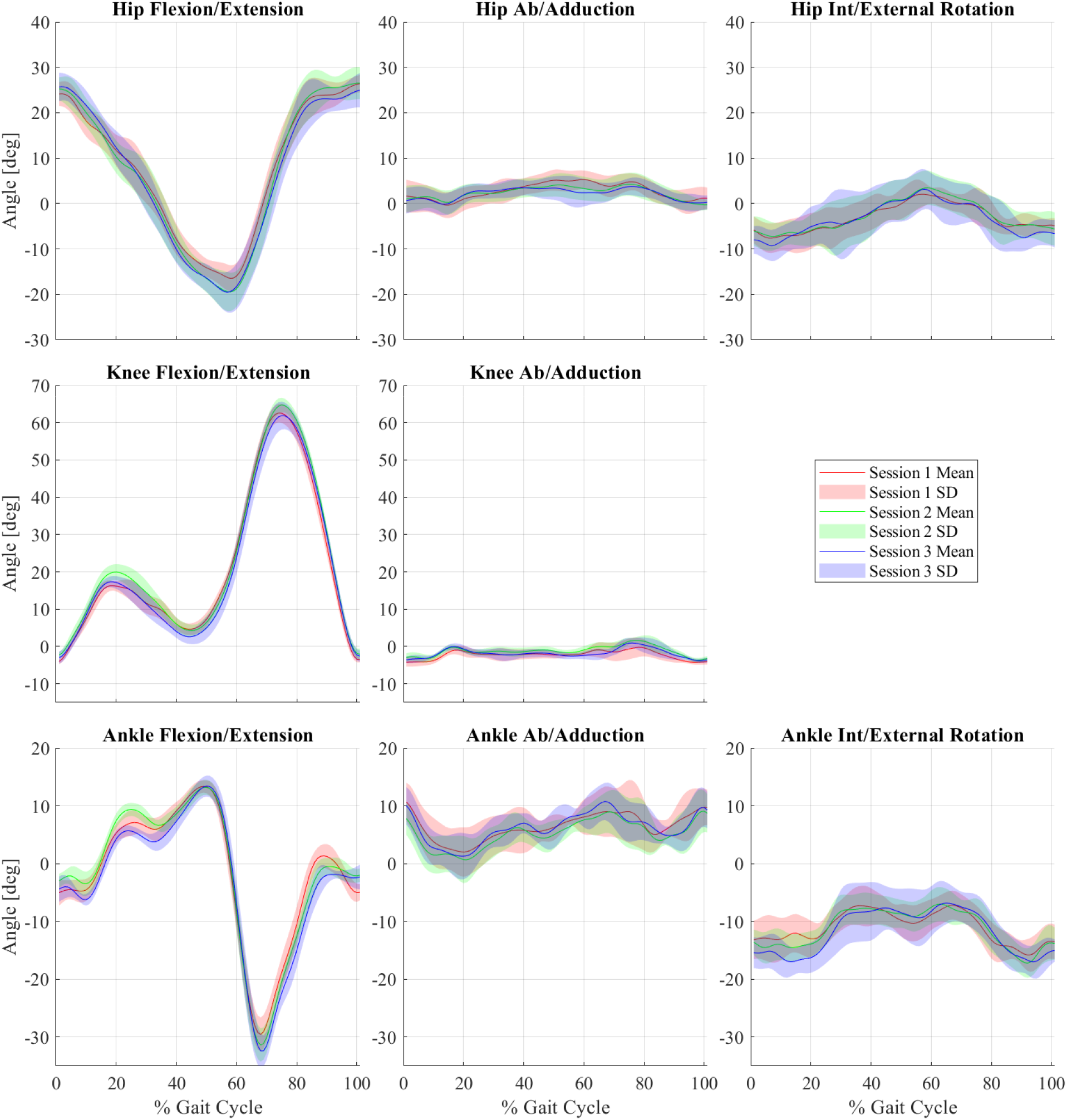
Lower limb joint angle patterns measured using the *Theia3D* IK model throughout the gait cycle, for all three sessions from one representative subject. Mean +/− SD for session 1 (red), session 2 (green), and session 3 (blue) are shown.

**Figure 3:**
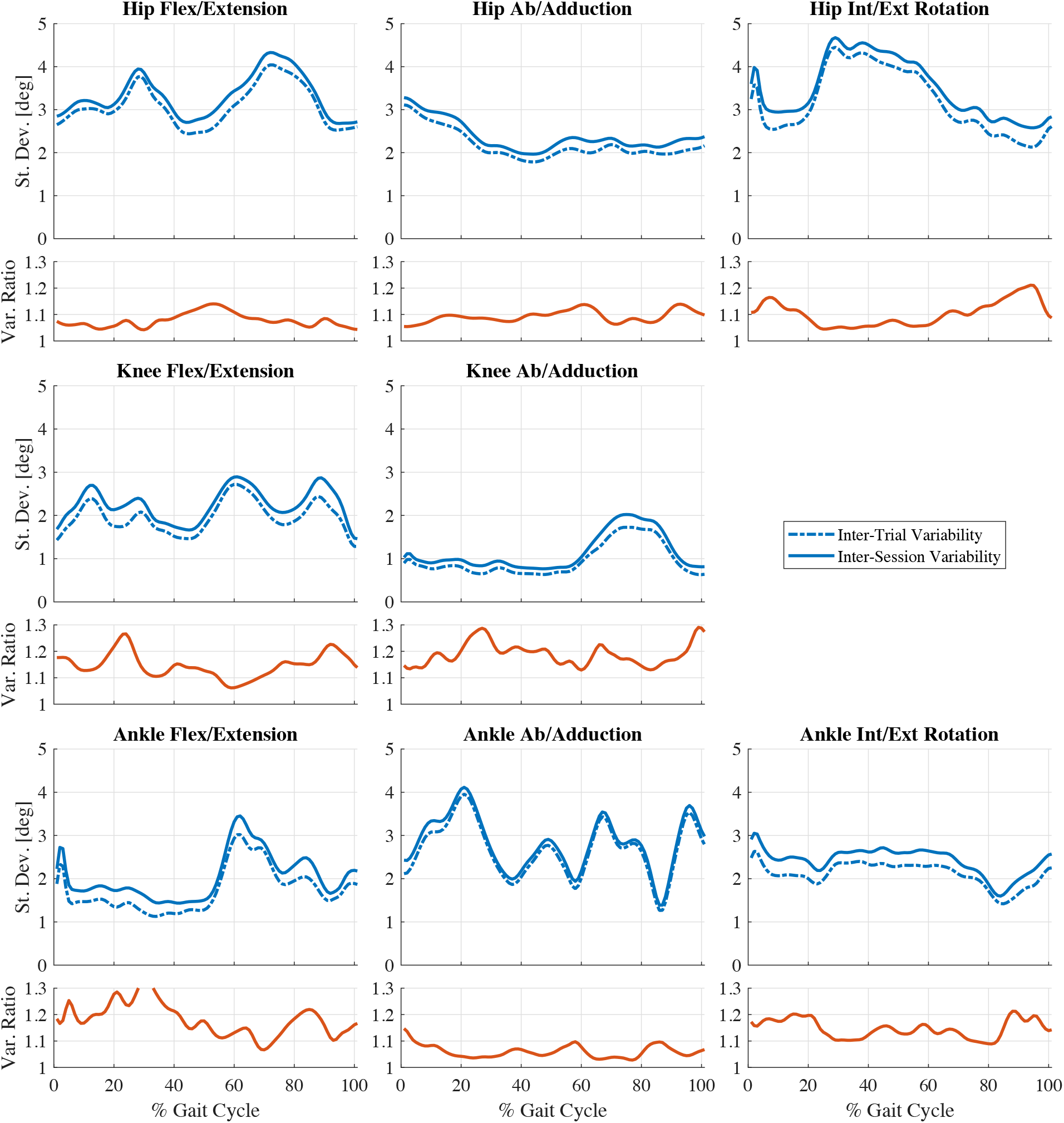
Average patterns of inter-trial variability (dashed lines) and inter-session variability (solid lines) expressed as standard deviations (St. Dev.) in degrees, plotted throughout the gait cycle for the lower limb angles of all subjects from the *Theia3D* IK model. The ratio of inter-session to inter-trial variability is included in the panel below each plot (Var. Ratio).

**Figure 4:**
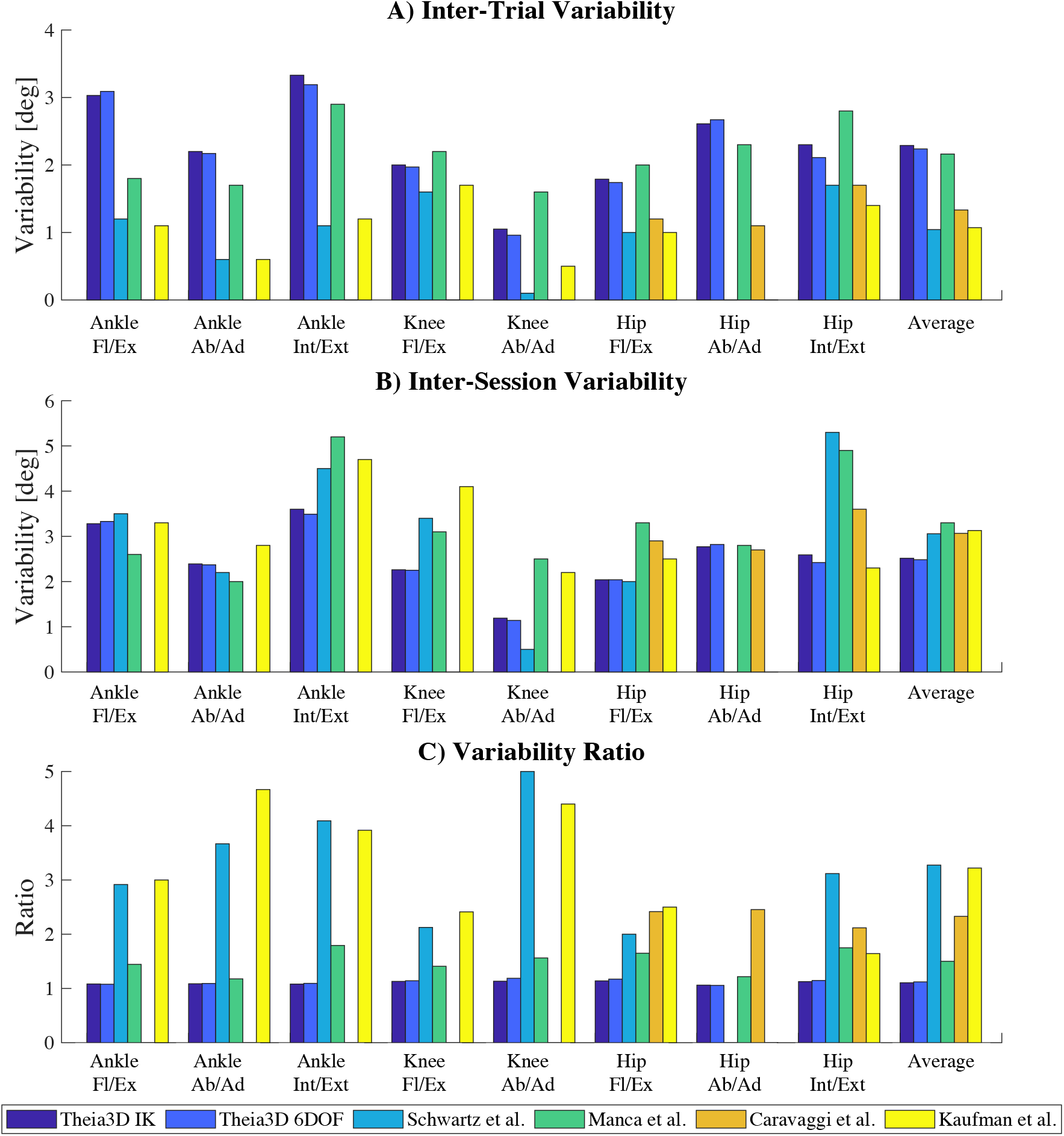
(A) Average inter-trial variability, (B) average inter-session variability, and (C) average variability ratio obtained in this study using the 6DOF model and IK model, and that from studies by Schwartz *et al.* (Schwartz et al., 2004), Manca *et al.* (Manca et al., 2010), Caravaggi *et al.* (Caravaggi et al., 2011), and Kaufman *et al.* (Kaufman et al., 2016).

The inter-trial and inter-session variability estimates were found to be very similar across the gait cycle for all measures, with the inter-session variability being larger than the inter-trial variability but with little difference between them (Figure 3). The inter-trial and inter-session variability measures were mostly below 4° except for some momentary peaks in hip flex/extension, hip int/external rotation, and ankle ab/adduction. The small differences between the inter-session and inter-trial variability are exhibited in the variability ratio, which is relatively constant and has an average of 1.1 or 1.2 for all measures (Figure 3, Figure 4C). Average inter-trial variation measures for the markerless kinematics were the largest among the compared studies for hip flex/extension, ab/adduction, int/external rotation, and ankle ab/adduction. The average inter-trial variability across all joints and planes was also the largest, at 2.2° (Figure 4A). Inter-session variability measures for the markerless kinematics were similar or lower than those from the included studies for all joint angles. Inter-session variation for hip int/external rotation, knee flex/extension and ab/adduction, and ankle flex/extension and int/external rotation angles were all the smallest by at least 0.5°, as was the average across all joints and planes, at 2.5° (Figure 4B). Average variability ratios for the markerless kinematics were the smallest among the included studies for all joint angles, with a maximum of 1.2 for knee ab/adduction and ankle flex/extension, and an average across all angles of 1.1 (Figure 4C). The average variability ratio of 1.1 indicates that performing multiple separate sessions increased the total variability of subjects’ measured kinematics by 10% on average.

## 4. Discussion

The use of *Theia3D* to calculate joint kinematics removed the reliance on an experienced examiner to identify anatomical landmarks and accurately place markers, substantially reduced the time to collect data, and resulted in kinematic data that was reliable between sessions on separate days.

Inter-trial variability was generally larger in this study (average = 2.2°) than those previously reported (1.0°-2.2°), however remained far below a suggested clinically acceptable threshold of 5° and subjects’ session mean waveforms were found to be consistent between sessions despite the greater intertrial variability. (McGinley et al., 2009; Caravaggi et al., 2011; Kaufman et al., 2016; Manca et al., 2010; Schwartz et al., 2004). The average inter-session variability measured in this study (2.5°) was the smallest across all other studies (3.1°-3.3°), indicating that measuring gait kinematics across multiple sessions using *Theia3D* introduces less variability compared to field-accepted marker-based motion capture systems. The variability ratios measured here were smaller than those in other studies for all examined joint angles, with the largest in the present study (1.2) being equal to the smallest reported value among the other studies (Manca et al., 2010). The small variability ratios obtained were due to the combined effect of larger inter-trial variability and smaller inter-session variability than those previously reported.

Repeatability measures obtained from the markerless kinematics were found to minimally differ when using the IK and 6DOF ankle models available within *Theia3D*, most notably for the ankle joint angles where the joint constraints differed between models. These findings indicate that the implementation of an IK approach to determine subject pose had little effect on the repeatability of the markerless kinematics, despite IK techniques having been shown in some studies to reduce the variability of kinematic measurements compared to direct kinematic (DK) approaches (Charlton et al., 2004; Kainz et al., 2017; Mantovani and Lamontagne, 2017). We believe instead that the greater repeatability of the markerless kinematics compared to those from previous studies is more likely due to the markerless tracking method itself. We acknowledge that greater differences may be seen if the knee and hip joints were additionally allowed 6DOF; however, other studies have shown similar reliability between IK and DK techniques and there is currently no implemented option for 6DOF knee and hip joints in *Theia3D* (Mentiplay and Clark, 2018; Horsak et al., 2018).

Given that our subject sample was made up of healthy adults with no neurological or musculoskeletal impairments, it is more likely that the greater inter-trial variability was a result of increased measurement noise within and between successive trials as opposed to greater subject gait variability. This higher level of inter-trial noise is likely caused by the markerless motion capture algorithm which uses a frame-by-frame approach to track subject movement. While this method has benefits such as not prescribing any expected movement patterns allowing it to track a wide variety of movements, it has the downside of potentially greater noise.

There are some limitations to this work that warrant consideration. We did not directly compare markerless and marker-based motion capture kinematics during the same trials because we wanted to perform the markerless data collections on unrestricted attire, and we have separately performed and reported on such a comparison. Despite being provided no instruction regarding attire, the participants wore mostly dark clothing during the data collection sessions, which is thought to provide a greater challenge in the accurate identification of anatomical features for the markerless system due to reduced contrast but has not been studied in depth. We also acknowledge that in order to perform lower limb kinematics, both legs must be visible and so this approach does have some limitations in terms of clothing, excluding, for example, full-length coats or skirts. Also, while markerless motion capture does not rely on skin-mounted markers and is therefore not affected by the inconsistent placement of markers between operators, it may instead be affected by systematic anatomical landmark calibration errors that could arise through the training of its neural networks. However, the accuracy of the anatomical labels within the training dataset cannot be benchmarked as there is no ground truth available for this data. Furthermore, while the markerless motion capture system is largely unrestricted with regards to the data collection environment, the data used in this study were collected in a laboratory space. Finally, the makeup of people included in the training images has not been fully documented and may contain biases with respect to various subject appearance characteristics. Thus, further work should be done to determine the sensitivity of gait kinematics to the collection environment, subject appearance and attire, and capture volume size, since these factors may differ in collected video data compared to the training dataset.

The findings presented here demonstrate that gait kinematics measured using *Theia3D* markerless motion capture are less affected by the use of multi-session protocols as indicated by the lower intersession variability and lower variability ratios compared to those previously reported for marker-based methods. The slight increase in inter-trial variability in combination with the decreased inter-session variability and ease of data collection is an acceptable compromise because of the potential for substantially more accessible data collections and more reliable longitudinal data.

## Conflict of Interest Statement

Scott Selbie is the President of Theia Markerless Inc. (Kingston, Ontario), the developers of *Theia3D.*

## Acknowledgements

This work was supported by an NSERC Canadian Graduate Scholarship (Master’s).

## Notes

### Competing Interest Statement

WSS is the President of Theia Markerless Inc. (Kingston, Ontario), the developers of Theia3D. He contributed to the conception and design of the study, critically revised the article for intellectual content, and provided final approval of the submitted version. WSS was not involved with the collection, analysis, or interpretation of data.

